# Electrolytic-microbubble dynamics delineate safety thresholds during intracortical microstimulation with flexible neural interfaces

**DOI:** 10.64898/2026.06.17.733032

**Authors:** Artem Iliasov, Haoran Ma, Fangyuan Li, Zhen Chen, Mingliang Xu, Chenhui Yu, Ruxin Li, Jinsong Wu, Fei He

## Abstract

Intracortical microstimulation (ICMS) with ultraflexible neural electrodes enables low-threshold, chronically stable, and high-resolution modulation of neural circuits, providing a promising strategy for sensory restoration and closed-loop neuromodulation. However, the microscopic mechanisms delineating its safe and effective current range remain unclear. Here, we combine intravital two-photon (2P) imaging and electrophysiology in awake mice to examine the current-dependent neurovascular outcomes of charge-balanced stimulation via ultraflexible arrays. We observed gas bubbles formed along the electrode during ICMS, with bubble size increasing quadratically with current amplitude, consistent with a Faradaic bubble-growth model. Intravital 2P imaging reveals that at low-to-moderate currents (20–40 µA), vascular leakage is small, spatially confined, and largely reversible, whereas higher currents (≥60 µA) induce a sharp transition to extensive, field-dominated extravasation and secondary vessel disruption. This transition coincides with immediate, stimulus-locked motor responses and the onset of electrode degradation. Multiphysics simulations reproduce the observed nonlinear leakage–current relationship by incorporating gas bubble–induced electric field redistribution and voltage-dependent vessel wall permeability. The model indicates that gas bubbles act as local electric-field modulators, concentrating suprathreshold fields near the bubble boundary at lower currents while shielding more distant vessel segments; at higher currents, this confinement breaks down and the system enters a field-dominated damage regime. Collectively, these findings define a mechanistically informed safety window for ICMS with flexible neural interfaces and identify bubble-assisted vascular permeabilization as a key failure mode at high currents, crucial for the design of future bidirectional brain-computer interfaces and high-precision neuroprosthetic protocols.

## INTRODUCTION

Intracortical microstimulation (ICMS) using invasive neural electrodes has emerged as a powerful tool in fundamental neuroscience and clinical applications (*1–5*). It offers micrometer- and sub-millisecond resolution in probing brain function, a level of precision that is unattainable through non-invasive methods. This remarkable spatial and temporal resolution enables the selective activation or inhibition of specific neuron populations within cortical layers, minimizing off-target effects and improving therapeutic outcomes (*6–8*). Moreover, ICMS facilitates the accurate mapping of functional brain areas, which contributes to a deeper understanding of sensory processing (*9*), motor control (*8*, *10*) and cognitive functions such as attention and memory (*11*). By directly delivering calibrated electrical pulses to the cortex, it can generate sensory percepts (*12*, *13*), including tactile feedback for prosthetic limbs (*7*), phosphenes for artificial vision (*9*, *14*, *15*) or auditory signals to restore hearing (*16*) in individuals with neurological impairments. Additionally, the therapeutic potential of ICMS extends to the treatment of conditions such as Parkinson’s disease, epilepsy, chronic pain and psychiatric disorders including depression, anxiety and addiction, through the precise targeting of the neural circuits involved in these disorders (*17*, *18*). In the future, next-generation devices will utilize continuous feedback to automatically adjust stimulation, a technique known as closed-loop neuromodulation or bidirectional brain-computer interfaces (BCIs), which marks the advent of adaptive and intelligent neuroscience (*19*).

Recently, ultrasmall and flexible neural electrodes—comparable in size to individual neurons—have transformed chronic neural recording and microstimulation (*20*, *21*). By matching the mechanical properties of brain tissue, they can integrate seamlessly with neural tissue, mitigate immune responses, and minimize glial scarring over months, making them ideal for central nervous system interfaces (*22–24*). In ICMS applications, they enable low-threshold, chronically stable neuromodulation at microscale with sub-millisecond precision (*25*). However, the biophysical effects of current delivery through such tiny electrodes remain poorly understood. For example, their flexibility and small size may complicate predictions of current spread and consistent electrode-tissue contact under dynamic conditions. Even moderate currents can generate high current densities and steep potential gradients at the electrode tip, which may potentially lead to micro-electrolysis, gas formation, and localized permeabilization of microvessels. In this microenvironment, strong local fields and Faradaic reactions may occur in single capillaries or extracellular spaces (*24*, *26*, *27*) giving rise to endothelial or neuronal membrane permeabilization (*28*), microbubble formation (*24*, *26*), and local heating (*29*)—effects that contribute to current-dependent nonlinearities that are undetectable by conventional bulk assays. Thus, a comprehensive reevaluation of the microscopic mechanisms that define safe and effective ICMS operation is imperative. Key questions remain about how localized electric fields interact with individual neurons and glial cells, how charge injection alters extracellular ion concentrations, and what thresholds cause tissue damage or unintended network activation (*1*, *30*, *31*).

While the safety assessment of neural stimulation has conventionally relied on empirical charge-density limits to evaluate reversible and irreversible tissue damage, these macroscopic metrics neglect the heterogeneous microenvironment surrounding ultrathin and flexible electrodes. For example, in ICMS it is still unclear how current amplitude and pulse dynamics, including duration, frequency, and waveform shape, influence local electrochemical, mechanical, and biological responses at the tissue-electrode interface (*18*, *26*, *32*). Excessive charge injection can initiate harmful reactions such as water hydrolysis, electrode corrosion, or the production of toxic reactive oxygen species, which may damage tissue and reduce the longevity of the device (*24*, *26*). The repeated expansion and contraction of electrode materials during stimulation can also induce mechanical stress, leading to microscale injury to neurons and glial cells (*22*, *23*). Biologically, unregulated parameters may trigger inflammation, neuronal death, or gliosis, which can degrade signal quality and device functionality (*18*, *22*, *23*). A more comprehensive understanding of these biophysical and cellular interactions is crucial for defining safe stimulation limits, enhancing the ICMS protocols, and designing more biocompatible, high-performance devices for long-term use. This will demand advancements in high-resolution imaging, computational modeling, and real-time electrophysiological monitoring.

Here, we combine intravital 2P imaging, extracellular electrophysiology, infrared behavioral tracking to systematically examine current-dependent neurovascular responses to ICMS in awake mice. By directly visualizing cortical microvasculature during stimulation, we identify distinct stimulation regimes that are not apparent from neural recordings alone. *In vivo* 2P imaging reveals that low-current ICMS produces little to no detectable vascular leakage, whereas intermediate currents give rise to small, spatially confined, and largely reversible extravasation events. At higher currents, ICMS induces gas bubble formation along the electrode and a sharp transition to extensive vascular disruption, including secondary vessel ruptures extending beyond the immediate electrode vicinity. Parallel electrophysiological recordings show that spiking activity and waveform features remain largely stable across tested current amplitudes. To interpret the observed nonlinear relationship between current amplitude, bubble formation, and vascular leakage, we employed multiphysics modeling as a complementary analytical tool. The model links the quasi-static electric field to voltage-dependent changes in vessel wall permeability and accounts for gas bubble–induced redistribution of the electric field. This framework reproduces the experimentally observed transition from localized to widespread leakage and provides a physical basis for the observed current thresholds. Our findings demonstrate that the effective operating range of ICMS is constrained not only by neuronal readouts, but also by microvascular electrochemical processes at the electrode–tissue interface. Rather than mechanical stress alone, bubble-assisted redistribution of the electric field emerges as a key factor governing the spatial extent and reversibility of vascular permeabilization.

## RESULTS

### ‘Four-in-one’ platform for multiscale analysis of ICMS

We developed an integrated multimodal platform (**Fig. 1**) to study how the local electric fields generated by the ultraflexible electrode interact with cortical microvasculature. As illustrated in **Fig. 1A**, the system combines four key modalities under a single cranial window: 1) 2P imaging of vasculature, electrodes, and gas bubbles; 2) ICMS; 3) extracellular recording via the same electrode; and 4) synchronized infrared behavioral monitoring in awake animals. We first optimized stimulation parameters *in vitro* using the fabricated 1.3-µm-thick ultraflexible platinum electrodes embedded in 1.2 % agarose, identifying pulse structures and current ranges that reliably produce visible gas bubbles under the optical microscope under controlled electrochemical conditions. For *in vivo* experiments, ultraflexible electrodes were implanted beneath a glass cranial window into the mouse somatosensory cortex (see **Materials and Methods**). Cortical microvasculature was labeled with 2-MDa FITC-dextran and visualized via 2P imaging during stimulation. This high-molecular-weight tracer was selected to conservatively detect vascular disruption: it does not extravasate through intact or mildly permeabilized endothelium, so observed leakage indicates either large-pore formation or structural vessel damage—not subtle BBB modulation. It also minimizes false positives and enables clear distinction between “no leak” and “leak” during threshold mapping.

**Figure 1.**
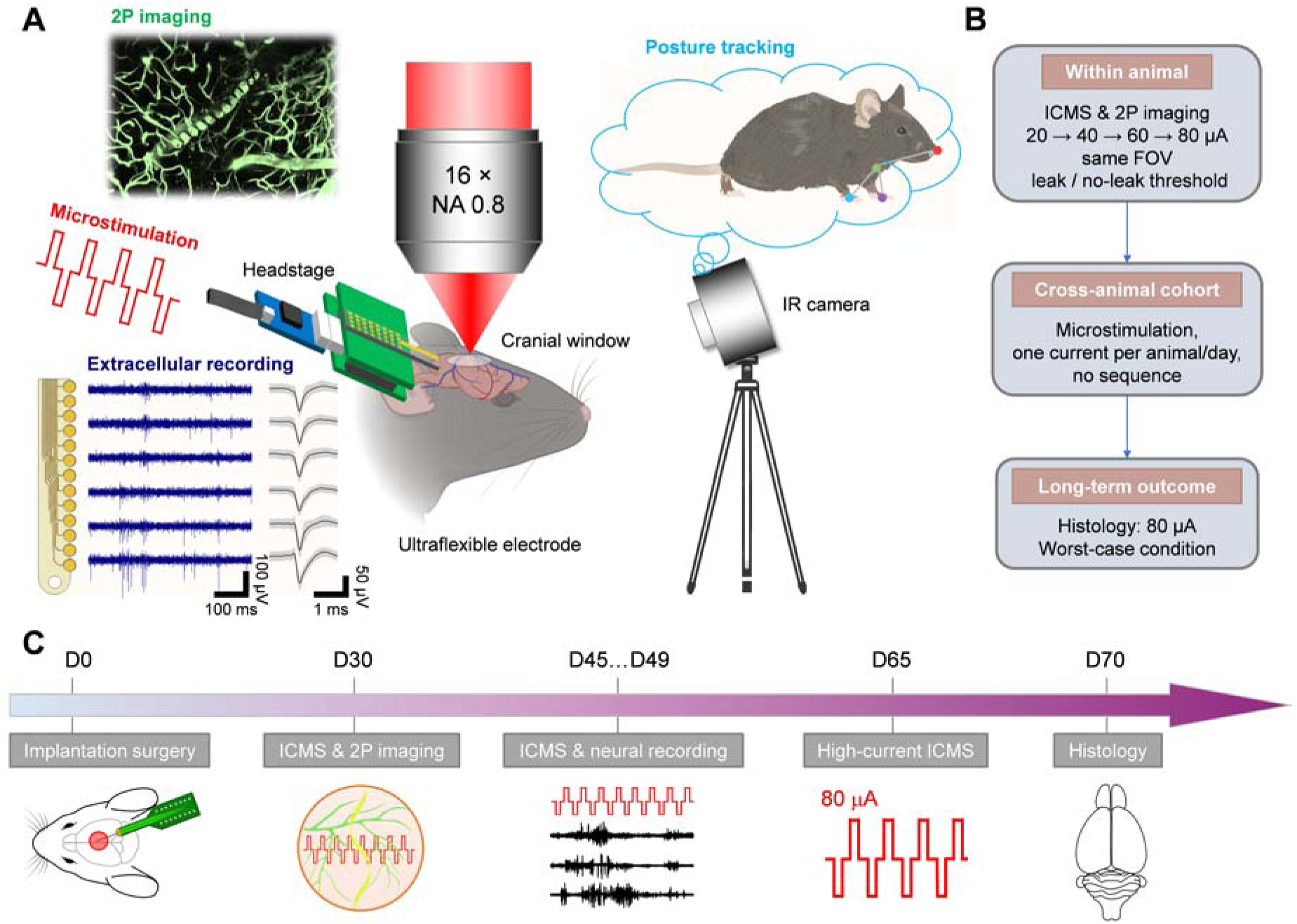
Integrated multimodal platform for assessing the safety window of ICMS in mice. **(A)** Schematic of **‘**four-in-one’ neural platform for awake mice including 2P imaging, extracellular recording, ICMS, and infrared behavioral tracking. **(B)** Overview of the experimental design: Within-session sequential ICMS (20–40–60–80 µA) mapped leak/no-leak thresholds in the same field of view (∼1 h). Daily ICMS sessions with varying amplitude (one amplitude per animal per day, delivered via distinct electrode channels) provided data for statistical analysis. Histology was performed only after 80 µA stimulation—at D70—as a delayed worst-case assessment. **(C)** Experimental procedures and timeline: After electrode implantation and cranial window surgery (D0), multimodal sessions began at D30. Intermediate electrophysiology and daily ICMS sessions with varying amplitude occurred at D45–D49. A final 80 µA ICMS at D65 probed worst-case outcomes, followed by terminal histology at D70.

The experimental design for mapping leakage thresholds and enabling statistical comparisons is outlined in **Fig. 1B**. In a within-animal sequential stimulation protocol, current amplitude was increased stepwise (20 → 40 → 60 → 80 µA) over ∼1 hour in total. At each amplitude, a 25 s stimulation epoch was delivered, followed by 10–15 min of time-resolved imaging to monitor vascular leakage in the same vessel segments before proceeding to the next amplitude. To avoid localized tissue conditioning, electrode polarization, or cumulative microenvironment alterations from the initial threshold mapping, subsequent daily sessions were performed using alternative, spatially distinct channels of the 32-channel array. After recovery, each mouse received only one stimulation amplitude daily (no escalation), enabling cohort-level statistical analysis of vascular leakage, electrophysiological stability, and behavioral responses. Histology was performed only after 80 µA stimulation—representing the worst-case condition—to assess delayed, chronic tissue-electrode effects. **Fig. 1C** shows the timeline: probe implantation, multimodal imaging, stimulation, and delayed histological assessment. This staged approach minimizes acute surgical confounds, allows neurovascular recovery and stabilization, and captures both immediate and delayed stimulation effects. Crucially, it enables direct cross-validation of electrochemical signals, vascular dynamics, neural activity, and behavior in the same subject—reducing animal use and boosting statistical power via co-registered measurements.

### Current-dependent gas bubble formation and electrode damage *in vitro*

Initially, we used an *in vitro* assay with a platinum electrode buried in agarose to screen stimulation parameters for efficient gas bubble generation and identify current ranges that preserve electrode integrity (**Fig. 2**). At 80 µA, time-lapse imaging (**Fig. 2A**) shows bubble nucleation at the active channel, progressive expansion during stimulation, attachment to the electrode surface, and gradual shrinkage post-stimulation. Bubble formation was always spatially confined to the stimulated site. At intermediate and high currents, multiple sub-visible microbubbles appeared along the electrode before rapidly coalescing into one dominant macroscopic bubble (**Movie S1**). We therefore quantified only this dominant bubble for comparison with *in vivo* 2P. To confirm the origin of the gas bubble formation, we monitored local chemical changes during stimulation in an *in vitro* gel. Electrical stimulation in FITC-containing gel induced a visible color change in the vicinity of the active contact site (**Fig. 2B, C**), which indicates a local pH shift caused by electrolysis. Furthermore, repeated stimulation caused a progressive decrease in open-circuit potential (OCP; **Fig. 2F**), which is consistent with acidification from water splitting. This aligns with the expected half-reactions: 2H_2_O → O_2_ + 4H^+^ (anode) and 2H_2_O + 2e^-^ → H_2_ + 2OH^-^ (cathode). A further calibration experiment demonstrated that the OCP correlates linearly with bulk pH (R^2^ = 0.99; **Fig. 2G**), which confirms that the OCP drift can reliably reflect the stimulation-induced pH changes.

**Figure 2.**
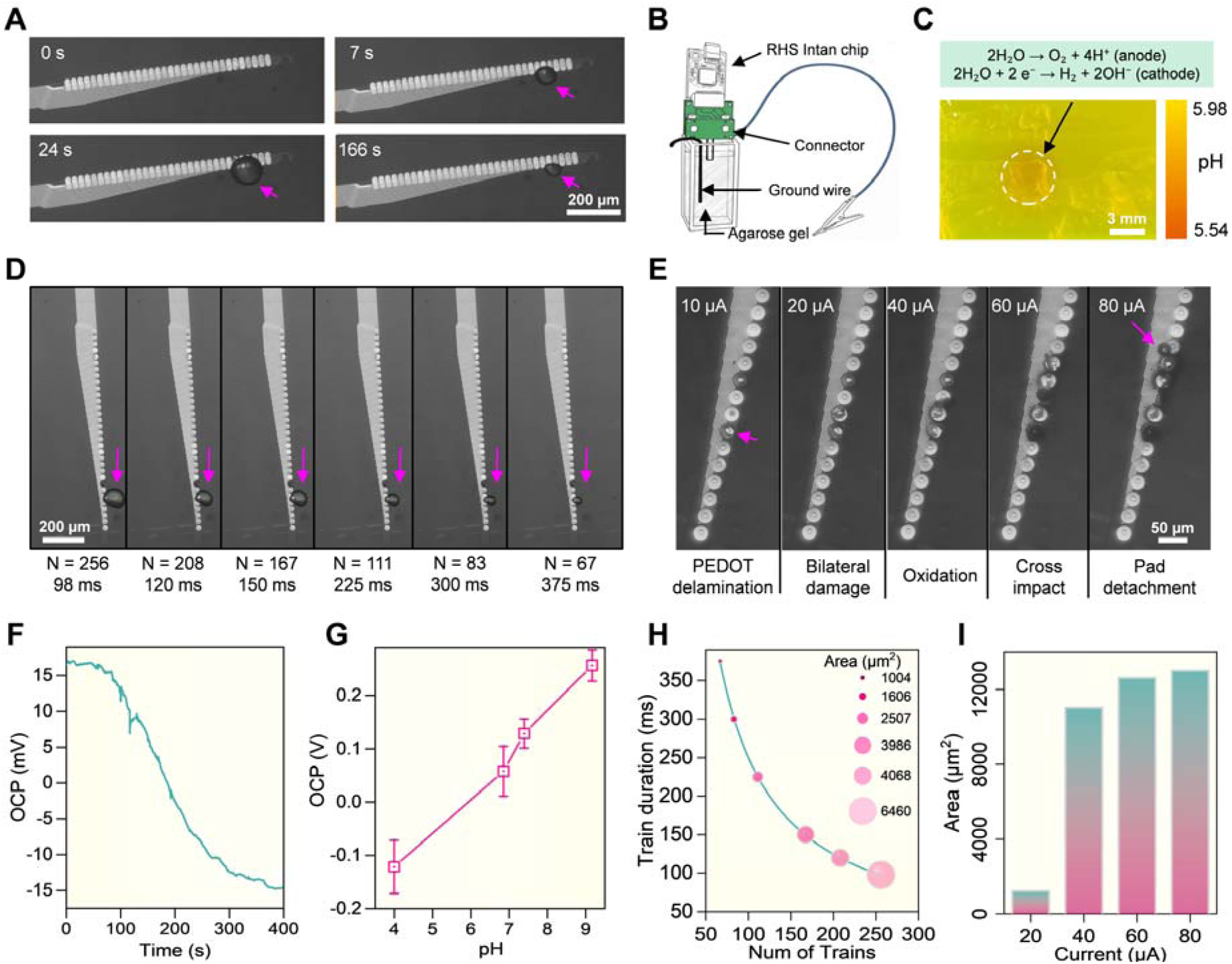
*In vitro* optimization of stimulation parameters for gas bubble generation and electrode integrity. **(A)** Time-lapse bright-field microscopy of gas bubble nucleation, growth, and dissolution on the electrode during and after stimulation. Representative frames: bubble nucleation and growth during the 25-s stimulation epoch (0–25 s), and subsequent dissolution after current cessation (166 s). Complete dissolution occurred at 257 s (not shown). **(B)** *In vitro* electrolysis setup for stimulation parameter screening: an ultraflexible microelectrode immersed in FITC-doped agarose gel (1.2 %, PBS buffered to pH 5.98 to maximize the dynamic range of fluorescence intensity), connected to stimulation/recording circuitry; a remote ground wire completed the circuit under optical access. **(C)** Localized electrochemical activity at the electrode–gel interface. The dashed circle marks the region where current injection induces a local pH change, visualized by the orange discoloration of the FITC-doped agarose gel due to water electrolysis. **(D)** Gas bubble formation under varying burst parameters: pulse train duration (T) and number of trains (N), at fixed frequency (66.7 Hz). Scale bar: 200 µm. **(E)** Effect of current amplitude (10–80 µA) on Pt electrode integrity. Representative micrographs show channel morphology post-stimulation. **(F)** Open-circuit potential (OCP) monitoring during three consecutive 25-s stimulation epochs. The shift in OCP reflects a progressive local pH decrease from ∼5.98 to ∼5.54 due to water electrolysis. **(G)** OCP–pH calibration curve (R² = 0.9909). Error bars: SD (n ≥ 3 independent trials). **(H)** Quantitative optimization of the stimulation burst structure. Gas bubble area (circle size) is plotted as a function of the number of trains (N) and pulse train duration (T). **(I)** Dependence of gas bubble size on current amplitude in 1.2 % agarose gel.

We then examined how burst structure affects gas bubble formation while maintaining a constant pulse frequency (66.7 Hz) and total number of delivered pulses. By varying the duration of each pulse train (T) and the number of trains (N), while keeping their product (N × T) constant, we redistributed identical total charge across different burst patterns (**Fig. 2D, H**). Both representative images and quantitative analysis showed that the area of the bubble strongly depended on the parameters of the stimulation burst: shorter T and larger N yielded more efficient bubble growth, with peak area at T = 98 ms and N = 256—the maximum supported by the hardware. This configuration was thus used in the subsequent experiments. Moreover, we examined how the current amplitude affects gas bubble formation and platinum electrode integrity (**Fig. 2E, I**). At a current range of 10-20 µA, the formation of bubbles was minimal and the electrodes remained intact. In the range 20-40 µA, localized bubbles were observed, yet the electrode structure was preserved. At ≥60 µA, the area of the bubbles surged and structural damage occurred, including cross-channel degradation and pad detachment. Thus, large bubbles and irreversible damage co-emerge when the current exceeds 60 µA. In contrast, at lower currents, controlled gas evolution can occur without interface failure. Meanwhile, gold contacts, whether bare or PEDOT:PSS-coated, exhibited notable damage and material loss at 20–40 µA (**Supplementary Fig. S1A-C**; **Movies S2** and **S3** demonstrate real-time PEDOT:PSS delamination and pad displacement). These results confirm the superior tolerance of platinum to high charge injection densities. It remains stable while gold interfaces degrade, even though those currents are considered to be safe for tissues. This is consistent with the established use of platinum in intracortical stimulation due to its higher charge-injection capacity and electrochemical stability (*26*). This *in-vitro* study demonstrates that gas bubbles are inherently formed during Faradaic reactions at microelectrodes, and the safe operating window for those ultrathin stimulation electrodes is narrow. It is jointly limited by gas evolution and electrode stability, which is defined here solely from electrochemical and materials perspectives.

### Current-dependent vascular leakage and bubble-mediated field confinement *in vivo*

All *in vivo* tests used awake adult C57BL/6J mice that were acclimated to head fixation (Materials and Methods). We implanted single-shank ultraflexible electrode arrays through a modified cranial window technique to record neural activity across the cortical depth. Experiments started at least 4 weeks after the surgery to allow for vascular recovery (*33*). We then performed *in vivo* 2P imaging to visualize cortical vascular response while delivering ICMS current via the implanted flexible neural interface (**Fig. 3**). The probe traversed the superficial vascular plexus at approximately 180–200 µm below the dura layer (**Fig. 3A, B**), exposing capillaries and small venules to the local electric field. Prior to stimulation (60 µA burst), fluorescence was confined to the vessel lumen, while after stimulation, a bright, diffuse FITC–dextran cloud extended far beyond the perivascular space, indicating large-scale, persistent leakage (**Fig. 3C, D**). While 2P imaging limits the apparent leakage extent to the field of view, widefield images under identical conditions (**Supplementary Fig. S2A, B**) show dextran extravasation spreading over hundreds of micrometers to millimeter scales at the ICMS currents of 60 µA or higher. Concurrently, leakage remained localized near the stimulated vessel.

**Figure 3.**
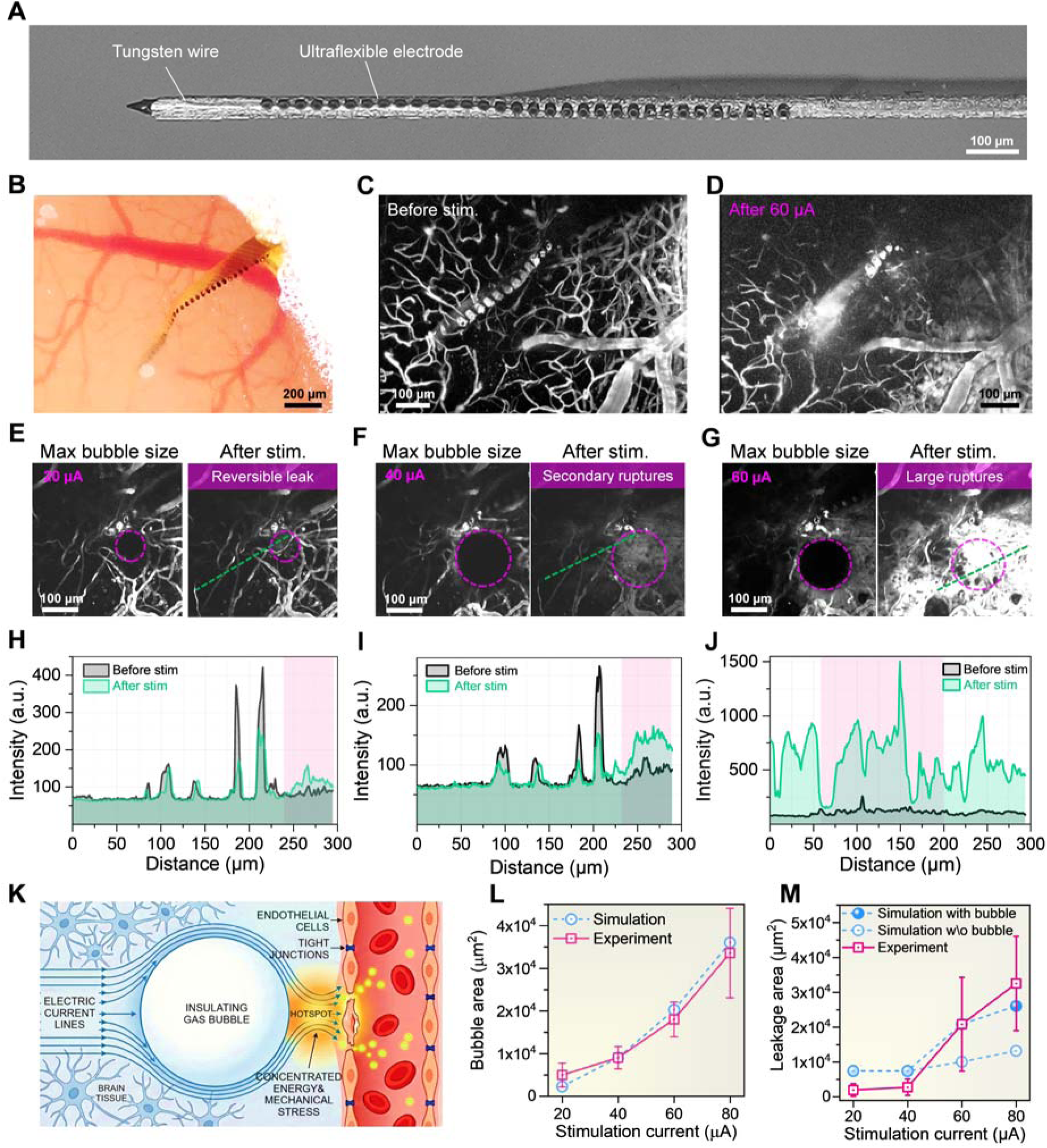
Current-dependent modes of vascular permeabilization revealed by *in vivo* 2P imaging and modeling. **(A)** Optical micrographs showing a single-shank probe assembled with sharpened tungsten wires. **(B)** Bright-field optical image of a cranial window with implanted flexible electrode in the mouse brain. **(C)** 2P images of cortical vessels before and **(D)** after 60 µA stimulation, showing FITC-dextran leakage. 3D time-lapse reconstructions are in **Movie S4** (pre-stim) and **S5** (post-stim). **(E-G)** Maximal gas bubble size (purple dashed circle) and corresponding FITC leakage patterns at (E) 20, (F) 40, and (G) 60 µA, respectively. **(H-J)** Measurement of the current-dependent leakage quantified via spatial intensity profiles. 1D FITC fluorescence profiles along the green line in (E-G); sharp peaks = vessel lumens, diffuse elevations = extravascular signal; pink shading = maximal bubble extent during stimulation. **(K)** Illustrative explanation of the proposed bubble-assisted blood–brain barrier (BBB) opening. **(L)** Relationship between the gas bubble area and stimulation current. **(M)** Relationship of the leakage area and stimulation current. Experimental data in (L) and (M): mean ± SEM (*n* = 7). A FEM model was used to simulate the microbubble generation in (L) and (M) during electrical stimulation (**Materials and Methods**), and the simulation results are in agreement with the experimental results only when the bubbles are considered in the simulation.

A conceptual framework is tentatively proposed to interpret the observed spatial and temporal patterns of vascular leakage, as depicted in **Fig. 3K**. The insulating gas bubble distorts the local current flow, causing the current lines to concentrate in the narrow gap between its surface and the adjacent capillary wall. This results in the creation of a localized hotspot with elevated electrical and mechanical stress. Crucially, the dissolution or collapse of the bubble abruptly releases this accumulated mechanical tension and localized electrical polarization, acting as the final trigger that transiently disrupts the endothelial tight junctions and increases the permeability, thereby allowing the leakage tracers to extravasate into the neural tissue. As a result, the leakage is restricted to the endothelium that is immediately adjacent to the bubble; whereas the vessel segments far from the bubble remain impermeable. Meanwhile, a portion of the injected charge drives the formation of gas, which reduces the effective electric field in the surrounding tissue. Thus, the bubble functions not only as a local field concentrator to focus the permeabilization at its interface, but also as a protective buffer to prevent the widespread vascular disruption, particularly near the threshold to open the blood brain barrier (BBB).

We then quantified the dependence of leakage on current amplitude, as shown in **Fig. 3H-J**. It was observed that at 20-40 µA (Regime I: Reversible), the leakage was spatially confined and transient. Specifically, at 20 µA, a small bubble formed beside one vessel segment, resulting in localized and rapidly fading leakage. At 40 µA, larger bubbles and occasional secondary leakage sites appeared, but the FITC tracer gradually washed out, which confirmed the reversibility. The intensity profiles of FITC exhibited narrow, localized signal increases, which were consistent with limited pore formation. At an ICMS current of ≥60 µA (Regime II: Catastrophic), large bubbles adhered to the vessels, and this was followed by sustained, high-intensity plumes spreading into the tissue. Intensity profiles revealed broad elevated plateaus, which indicated the rupture of extended vessel segments. Notably, the erythrocyte extravasation (identified by particle size and the absence of FITC signal; **Movie S6**, **Fig. S2C, D**) occurred at 60-80 µA, providing direct *in vivo* evidence of structural disruption of the vessel wall. The onset of leakage coincided with the collapse of the bubble rather than its growth, and it reached its peak while the bubble was partially present. Subsequently, it declined gradually over 20–30 minutes through diffusion, rather than rapid resealing (**Movie S7**). To investigate whether the bubble-mediated field redistribution accounts for the current-dependent bubble size and leakage area, a COMSOL finite-element model was constructed (**Materials and Methods**). The tissue was modeled as a conductive medium surrounding a circular electrode, and bubble growth was modeled using an ordinary differential equation (ODE) based on Faraday’s law and Laplace pressure. The results in **Fig. 3L** indicate that the simulated steady-state bubble area increased almost quadratically with current and closely matched the experimental data.

To further model the leakage, we introduced a nearby vessel segment with wall permeability dependent on local transmembrane voltage. A phenomenological bubble effect was incorporated via a size-dependent scaling factor that modulates the effective transmembrane voltage. It was found that at low currents, the model predicted only transient, limited permeability increases, which is consistent with reversible leakage. As current increased, the local electric field exceeded a critical threshold, causing permeability to fail to recover rapidly and resulting in sustained tracer extravasation. The simulated leakage–current relationship (**Fig. 3M**) matched the experimental observation: a gradual increase at low currents, followed by a sharp rise at intermediate currents—marking the onset of persistent leakage. Although the model slightly underestimated the abruptness of this transition (likely due to omitting continuous blood flow), it captured the qualitative shift from reversible to sustained barrier disruption using a single parameter set that also fit the bubble-size data. These results indicate that the redistribution of the electric field induced by bubbles, combined with the voltage-dependent endothelial permeability, is sufficient to explain the observed leakage transition.

In the experiment, we observed that bubble formation and vascular leakage varied across different sessions, which reflected the sensitivity to the local electrode–vessel geometry. However, leakage did not occur in every session, even when the current was at 60–80 µA. Furthermore, in some sessions, bubble was not detected during stimulation. We report the per-session outcomes (leak / no leak / no bubble) in **Supplementary Fig. S1D, E**. Time-resolved imaging reveals this variability within single sessions: repeated stimulation of the same electrode channel can sequentially produce multiple bubbles, but leakage was not observed (**Movie S8**). It may explain the large inter-animal variation in the leakage area (**Fig.3M**).

### Stable unit tracking and electrode performance after repeated ICMS

To test whether the stimulation-induced vascular leakage affects neuronal activity or electrode performance, we recorded extracellular signals from the same 85 channels in 6 mice for four consecutive days (∼540 total hours; **Materials and Methods**), as shown in **Fig. 4**. In each daily experiment, 30-minute electrophysiological recordings were performed on head-fixed animals before ICMS, followed by a single 25 s stimulation epoch delivered at a given current amplitude (20, 40, 60, or 80 µA). After a 4–5 min recovery period, post-stimulation activity was recorded for 1 hour. Each animal was exposed to only one stimulation amplitude per day. To evaluate the functional longevity of the neural interface, we monitored recording quality and electrode integrity during chronic microstimulation. Longitudinal tracking of individual units (one representative subject) showed that median firing rates (FR) remained stable across four days of increasing stimulation intensities from 20 to 80 µA (**Fig. 4A**). However, while firing patterns were preserved, we observed channel-specific fluctuations in spike amplitudes (**Fig. 4B**). These changes were closely associated with shifts in electrode impedance (**Fig. 4C**). Notably, on the stimulated channel (No. 30), current injection at 60 µA caused a rapid impedance drop and convergence with a neighboring channel (No. 7), indicating an inter-channel short circuit due to insulation failure (**Fig. 4C**).

**Figure 4.**
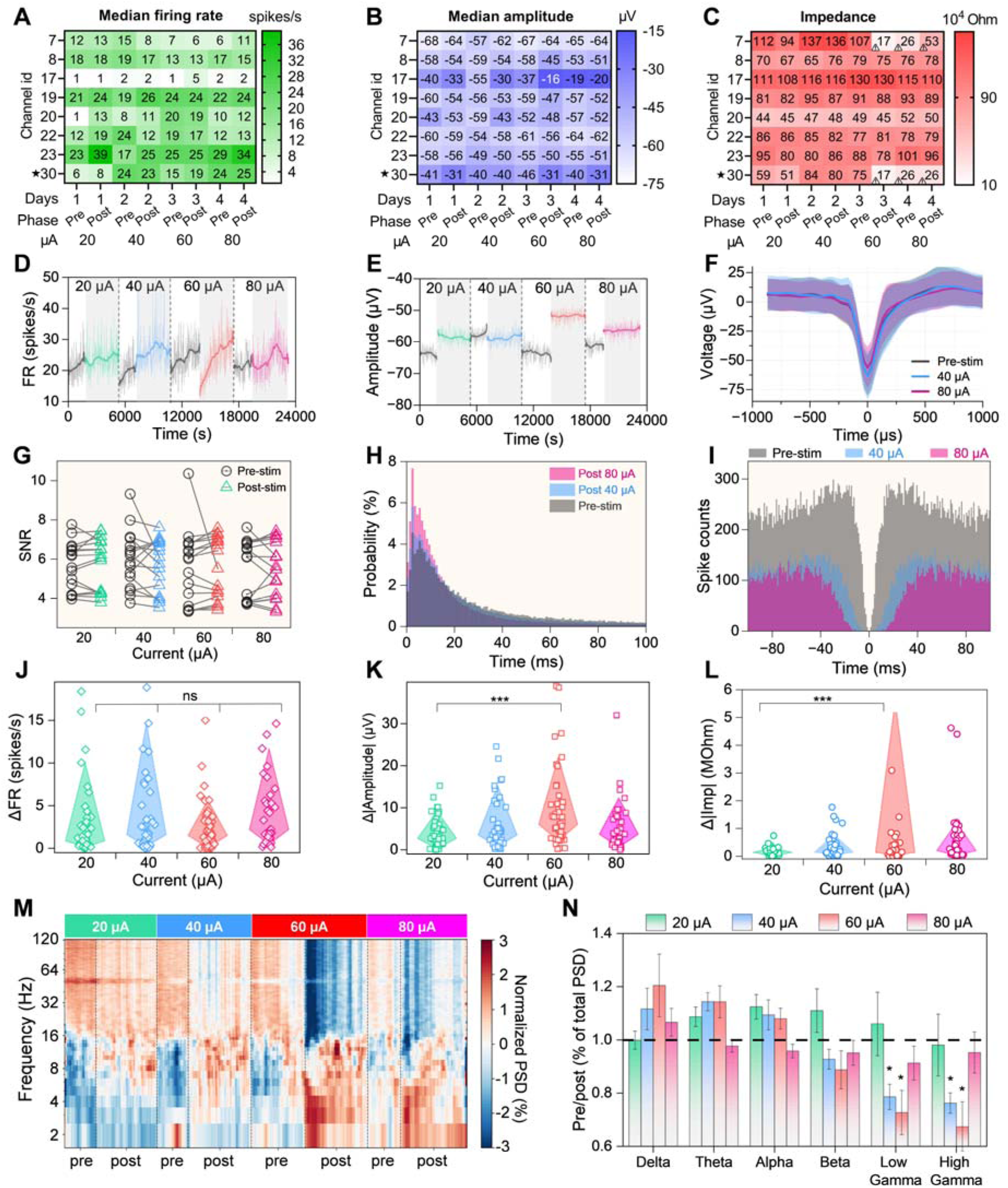
*In vivo* evaluation of the impact of ICMS on neural activity and electrode performance. **(A–C)** Heatmaps representing longitudinal tracking of median firing rate, spike amplitude, and electrode impedance for 8 neural units (one representative subject) across 4 days with increasing stimulation intensities 20—80 µA. Channel 30 (asterisk) denotes the stimulated electrode; convergence with channel 7 indicates insulation failure at 60 µA. **(D, E)** Representative time courses for a single channel. **(F, H, I)** Stability of a single unit (Channel 23) shown via average spike waveforms, ISI distribution, and autocorrelograms (ACG) (see **Supplementary Fig. S3** for data from additional isolated units). **(G)** Population-level SNR changes. **(J–L)** Statistical analysis of absolute changes (|Δ|) across all subjects and units. All groups showed significant deviation from zero (one-sample *t*-test, *p* < 0.001, not shown). Brackets indicate inter-group comparisons via one-way ANOVA; ***p < 0.001, ns: non-significant. **(M)** Time–frequency representation of normalized LFP power (PSD, % total power) across stimulation conditions (20–80 µA). **(N)** Quantification of band-specific power (pre/post, normalized to baseline) across canonical frequency bands. Gamma-band activity exhibits a reduction following stimulation, while lower-frequency bands remain relatively stable. Asterisks indicate significant differences (*p < 0.05).

Despite stimulation-induced changes in spike amplitude and impedance, the overall waveform shape remained largely preserved across conditions (**Fig. 4F**). Average spike waveforms before and after stimulation showed minimal distortion, with only modest reductions in peak-to-trough amplitude at higher current levels. Consistently, signal-to-noise ratio (SNR) remained stable across all stimulation intensities (**Fig. 4G**), indicating that unit isolation quality was maintained throughout the experiments. Analysis of spike timing statistics revealed subtle but consistent changes in firing dynamics. Inter-spike interval (ISI) distributions exhibited an increased probability of short intervals following stimulation, particularly at higher amplitudes (**Fig. 4H**). In parallel, autocorrelograms (ACG) showed elevated spike counts at short time lags (10–50 ms), indicating enhanced temporal clustering of spikes (**Fig. 4I**). Importantly, the refractory period was preserved across all conditions, supporting the conclusion that these effects do not arise from recording artifacts or unit contamination.

Population-level analysis across all subjects and units revealed that microstimulation induced measurable shifts in all parameters. One-sample t-tests against a null hypothesis of zero change confirmed significant effects (p < 0.001) for ΔFR, Δ|Amplitude|, and Δ|Impedance| across all intensities (**Fig. 4J–L**). While ΔFR remained statistically uniform between the 20–80 µA groups (**Fig. 4J**, one-way ANOVA), the amplitude and impedance changes were intensity-dependent. The absolute change in amplitude was significantly higher at 60 µA compared to 20 µA (**Fig. 4K**, p < 0.001), mirroring a sharp increase in impedance shifts at the same threshold (**Fig. 4L**). The lack of further statistical divergence at 80 µA suggests that the primary physical alterations to the electrode–tissue interface, such as insulation delamination, are largely initiated at the 60 µA level.

To assess how microstimulation affects local field potential (LFP) activity, we analyzed band-specific power dynamics across the same stimulation paradigm. Time–frequency analysis revealed that LFP power remained largely stable across low-frequency bands (delta, theta, and alpha), with only minor fluctuations before and after stimulation (**Fig. 4M**). In contrast, higher-frequency components exhibited more pronounced changes. Quantification of band-specific power showed that gamma-band activity was selectively reduced following stimulation (**Fig. 4N**). This effect was most evident in the low and high gamma ranges, where post-stimulation power decreased relative to baseline, particularly at intermediate and higher stimulation amplitudes (40–60 µA). In contrast, lower-frequency bands did not show consistent or significant deviations from baseline levels. Temporal analysis of individual frequency bands confirmed these trends. While delta, theta, and alpha bands remained relatively stable over time (**Supplementary Fig. S4A-C**), beta and gamma bands displayed transient reductions in power following stimulation epochs (**Supplementary Fig. S4D-F**). These changes were reproducible across animals and stimulation days.

### Behavioral responses and histological tissue-electrode interface integrity after ICMS

To assess stimulation-evoked behaviors, we simultaneously recorded infrared videos of head-fixed mice, in parallel with 2P imaging, electrophysiology and stimulation, as shown in **Fig. 5A, B**. The camera was focused on the muzzle and forepaws of the animal, and the animal body was unrestrained within the cylindrical holder, which allowed for the detection of rapid, stimulus-locked movements of the forepaws and body. **Fig. 5B** shows representative infrared frames captured before, during, and immediately after stimulation, demonstrating the abrupt motor response to suprathreshold stimulation. We quantified the instantaneous velocity (px/frame) using DeepLabCut across three periods: pre-stimulation, 25-second stimulation, and post-stimulation, respectively (**Fig. 5C-E**). Quantitative analysis revealed that low-amplitude stimulation (20 µA) did not evoke any detectable behavioral changes. The mice remained immobile, with only occasional spontaneous grooming that was indistinguishable from the baseline. In contrast, stimulation at current ≥40 µA triggered immediate, stimulus-locked motor responses, including vigorous forepaw movements, attempts to escape the holder, and occasionally vocalization. All these behavioral responses ceased promptly after the stimulation ended and the mice then either briefly groomed or returned to immobility.

**Figure 5.**
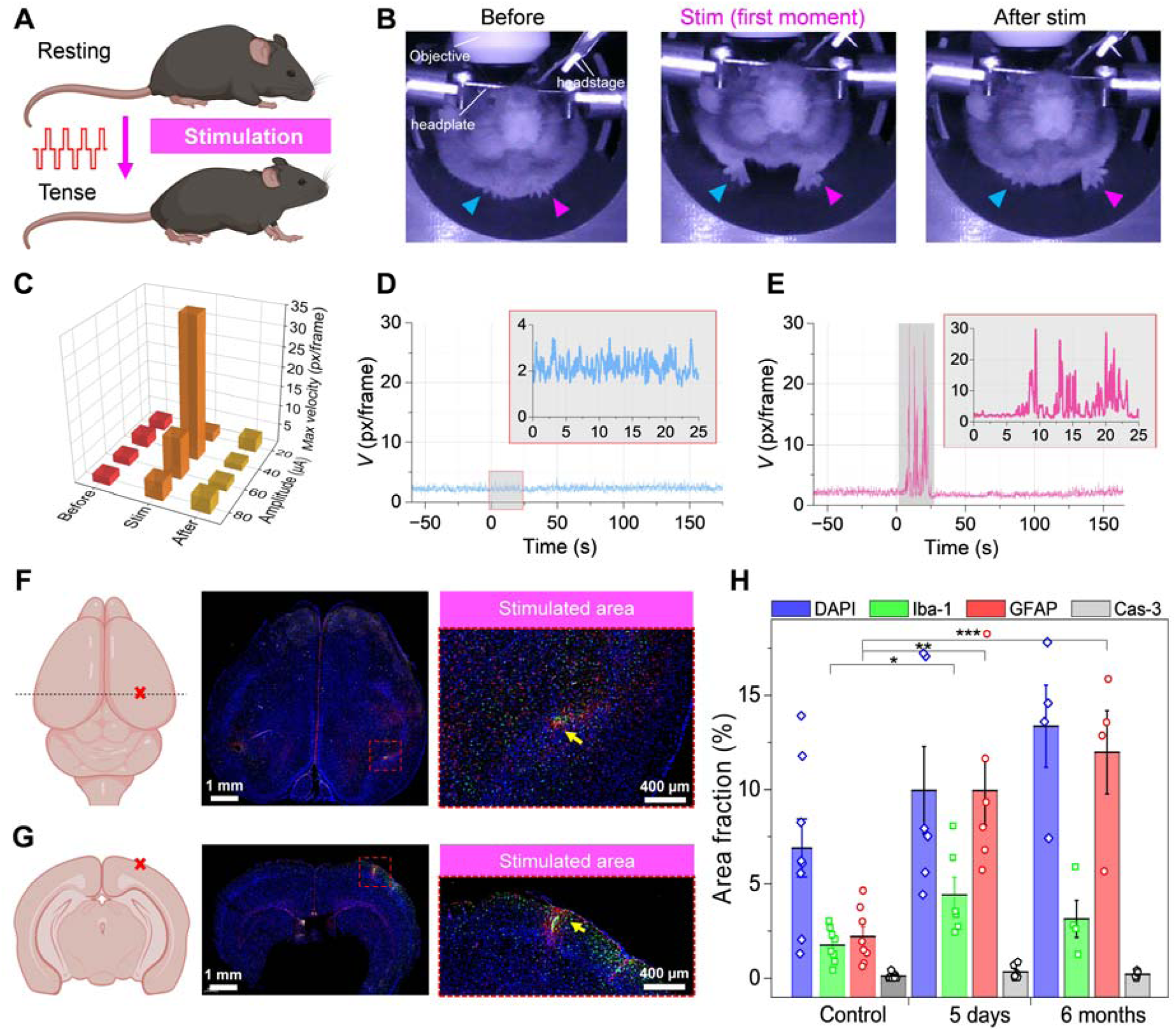
Behavioral and histological assessment of stimulation impact. (**A**) Schematic representation of stimulation effect on the mouse’s posture (resting vs. tense). (**B**) Representation IR images of the stimulation-evoked motor responses on a head-fixed mouse. (**C**) 3D heatmap of peak movement velocity (px/frame) across current amplitudes and time phases (Before, Stim, After). (**D, E**) Velocity time courses for control (D) and 60 µA stimulation (E), respectively. The red shading indicates the course under a 25-s stimulation train. **(F, G)** Histological validation of the electrode implantation sites. **(F)** Left: Schematic of the whole mouse brain (dorsal view) indicating the coronal/horizontal section plane (dashed line) and the stimulation site (red cross). Middle: Representative whole-slice immunofluorescent image. Right: Magnified view of the stimulated area (red dashed box), with the implantation site indicated by the yellow arrow. **(G)** Left: Schematic of a coronal brain cross-section indicating the cortical stimulation site (red cross). Middle: Representative immunofluorescent coronal slice. Right: Magnified view of the stimulated area with the implantation site indicated by the yellow arrow. Immunofluorescent staining markers: GFAP (astrocytes, red), Iba1 (microglia, green), and DAPI (nuclei, blue). (**H**) Time course of inflammatory/apoptotic marker area fractions in stimulated vs. contralateral regions (Control, 5 days, 6 months post-stimulation). *p < 0.05, **p < 0.01, ***p < 0.001 vs Control.

We further evaluated microglial and astrocytic activation through histology after last 80 µA stimulation. Representative horizontal and coronal slices are shown in **Fig. 5F, G**. We evaluated the inflammatory response and potential tissue damage through immunohistochemical analysis at two time points: 5 days and 6 months post-stimulation (**Fig. 5H**). Quantitative analysis of area fractions for GFAP (astrocytes), Iba-1 (microglia), and cleaved caspase-3 (apoptosis) revealed a spatially confined and time-dependent glial response. Microglial activation, marked by Iba-1 expression, showed a significant initial surge at 5 days post-stimulation (\**p* < 0.05 compared to control). However, by the 6-month mark, Iba-1 levels declined to the point where they showed no statistically significant difference from the contralateral control. In contrast, astrocytic activation exhibited a more persistent and progressive trend; GFAP levels rose significantly at 5 days (\*\**p* < 0.01) and showed an even more pronounced increase at 6 months (\*\*\**p* < 0.001), indicating the maturation of a stable glial scar. Importantly, despite the high-current stimulation (80 µA), the total cell density (DAPI) remained stable across all groups with no statistically significant differences. Furthermore, cleaved caspase-3 levels were negligible and remained at near-zero baseline levels at both time points (**Fig. 5H**). This absence of apoptotic signaling, combined with the lack of tissue voids or structural disruption, confirms that the stimulation protocol does not induce mass cell death or irreversible neurotoxicity, even in the long term.

At 6 months, **Supplementary Fig. S5** reveals a dense GFAP-positive astrocytic encapsulation surrounding the implant site, indicative of a mature, stable glial scar. In contrast to the acute phase, Iba1-positive microglial expression in the stimulated area has subsided, appearing comparable to the levels observed in the contralateral cortex. Furthermore, cleaved caspase-3 staining remains negligible, confirming the absence of chronic neurotoxicity or ongoing apoptosis following the stimulation protocol.

## DISCUSSION

Our study challenges the prevailing view in previous ICMS research that electrochemical gas formation is merely an undesirable side effect or a passive marker of overstimulation (*2*). By combining electrophysiology with high-resolution 2P vascular imaging, we demonstrate that electrochemically generated microbubbles act as dynamic biophysical modulators at the electrode–tissue interface. Rather than inducing immediate failure, the emergence of the gas phase establishes a clear two-regime threshold. At low-to-moderate currents, microbubbles serve as localized “electrical lenses” that focus the electric field into a narrow ring at the bubble boundary. This field redistribution is predicted to lower the effective transmembrane voltage required to disrupt endothelial barrier function, enabling reversible leakage of very large macromolecules (2 MDa) at currents well below those causing widespread tissue damage in prior field-dominated studies. Conversely, at higher currents, the combination of bulk field escalation and rapid bubble expansion drives the system into a catastrophic damage regime. Thus, understanding these specific bubble thresholds transforms gas evolution from an unpredictable hazard into a controllable phenomenon that may be leveraged for spatially selective vascular permeabilization. Multiphysics modeling confirms this mechanism, accurately reproducing the nonlinear dependence of bubble size and leakage area on current while validating the underlying field redistribution.

These findings provide mechanistic context for classical safety benchmarks (Shannon and McCreery) and identify the microscale events preceding macroscopic tissue injury (*34*). Our stimulation parameters, plotted on a Shannon diagram, span the critical transition from safe operation to damage. Behavioral and histological data support and refine the vascular observations. At low currents (∼20 µA), stimulation induces weak or absent motor responses and small, localized, largely reversible vascular leakage, consistent with near-threshold effects. At intermediate currents (∼40 µA), behavioral responses become robust while vascular integrity is largely preserved, defining a transitional regime. At higher currents (≥60 µA), irreversible failure emerges, characterized by vascular damage and electrode degradation. Collectively, these results define a mechanistically grounded operating window for ICMS. Classical models accurately predict macroscopic risk at high charge densities but fail to capture the key driver of the transition. Our results reveal a clear two-regime mechanism governed by the balance between electrochemical gas evolution and bulk electric fields. Thus, our findings suggest that gas evolution may represent an early physical event preceding Shannon-type stimulation damage. Monitoring of bubble thresholds provides a practical route to enhance safety protocols for the flexible neural interfaces, which enables real-time *in vivo* visualization and multimodal readout.

This mechanistic insight reframes gas formation from a safety hazard into a tunable biophysical process—distinct from recent studies on electric-field–driven microglial and BBB responses under conventional ICMS (*35*). Those studies showed field-dependent modulation using rigid electrodes at low duty cycles but did not account for the cavitation events we observe. Using ultrathin electrodes and real-time imaging, we demonstrate that bubble formation distinguishes field-dominated disruption from bubble-assisted vascular permeabilization. Conceptually, this mechanism resembles focused ultrasound (FUS)-induced BBB opening, where intravascular microbubbles—driven by acoustic pressure—cause transient permeability. In FUS, safety and efficacy depend critically on acoustic thresholds: Mechanical Index ∼0.4–0.5 and peak negative pressure ∼0.2–0.5 MPa; exceeding these increases hemorrhage and vascular damage risk (*36*). In both approaches, BBB opening can be spatially confined, reversible, and mechanically coupled to bubble dynamics. Crucially, bubble location differs: FUS microbubbles circulate intravascularly, whereas electrolytic bubbles form and remain anchored at the electrode–tissue interface—outside the vessel lumen—avoiding convective transport. Thus, embolic mechanisms are negligible; instead, local dielectric redistribution and bubble-associated mechanical stresses are likely the primary processes governing permeability changes. While the present study cannot fully separate these effects, the strong correspondence between modeled field amplification and experimentally observed permeability thresholds suggests that electrical lensing may act as an initiating mechanism of vascular permeabilization.

Prior work on electrical BBB permeabilization has primarily framed safety and efficacy in terms of electric field amplitude, pulse duration, and pulse number, largely independent of electrochemical phenomena. *In vitro* and microfluidic models have established that reversible opening is achievable within a narrow window (∼400–600 V/cm), but these platforms typically assume homogeneous tissue conductivity and ignore gas-phase formation (*28*). Our work reveals electrochemically generated gas bubbles as a critical driver of vascular permeabilization. Unlike high-voltage, whole-brain protocols using kilovolt pulses, our approach targets single vessels with ultralocal precision (*37–39*). Here, the initial onset of targeted permeability is not driven by bulk field escalation but by the emergence of a bubble-assisted microphysical state. This introduces an additional control dimension absent from microfluidic models: in the intact cerebral microvasculature, reversibility is governed not only by electric field magnitude but also by electrode-driven gas formation and collapse (*40*). This implies that *in vivo* safety boundaries cannot be fully captured by field-based metrics alone.

The ability to control this bubble-assisted regime has direct implications for molecular delivery. However, a fundamental distinction must be made between cellular electroporation—the transient poration of individual lipid bilayers widely characterized in literature (*41–43*) — and the vascular permeabilization observed in our study. While prior reports focus on optimizing field parameters for transmembrane transport, they often bypass the mechanical and electrochemical stressors acting on the integrated neurovascular unit. Only isolated in vitro studies have begun to address how electrical fields perturb endothelial barrier function, yet these lack the complex in vivo environment where gas evolution occurs (*44*). Our work bridges this gap, showing that bubble-mediated ‘electrical lensing’ creates a regime that transcends simple membrane poration. Consequently, the biophysical limits of electrode-mediated delivery remain insufficiently characterized.

Here, we address this threshold by mapping the transition from reversible vascular permeabilization to disruption in vivo. Unlike prior studies focused on delivery efficiency or endpoint histology, our real-time imaging shows that vascular permeability is tightly linked to short-lived bubble dynamics. This reveals a mechanistically grounded safety window—balancing microvascular integrity and electrochemical side effects. Gas evolution—long theorized but rarely documented in vivo—is thereby transformed from an unquantified hazard into a predictable boundary condition. Depending on the application, this boundary can be avoided (to preserve vascular integrity) or engaged (to enable enhanced permeability for targeted delivery). Critically, the high-intensity fields used here differ fundamentally from physiological signaling; recent work shows gap-junction–mediated electrical coupling between endothelial cells drives rapid vasodilatory responses *in vivo* (*45*). However, while intrinsic endothelial coupling enables adaptive blood flow regulation, our results show that externally applied electric fields can drive the vascular system into non-physiological electrochemical states—where classical neurovascular principles no longer hold. Specifically, field-induced gas evolution and cavitation—processes tightly linked to vascular disruption—are mechanical and chemical stressors fundamentally distinct from endogenous signaling.

Several limitations of this study should be acknowledged. First, experiments were performed in awake, head-fixed mice; while this allowed direct observation of physiological responses, species-specific differences in vascular architecture and tissue conductivity limit direct extrapolation to humans. Second, our results are specific to ultrathin, flexible electrodes. Such probes produce highly localized electric fields and elevated current densities near the surface, conditions that differ from conventional rigid microelectrodes. Third, measurements were restricted to the superficial cortical microvasculature; thresholds may differ in deeper brain regions or white matter where vascular geometry varies. Finally, although acute histology did not reveal gross necrosis, this study was not designed to assess cumulative inflammatory changes following chronic stimulation over months.

Our findings support shifting from static, charge-based safety thresholds to dynamic, mechanism-driven safety frameworks. Gas bubble formation, transient impedance changes, and vascular permeability are tightly linked—providing a practical foundation for bidirectional neural interfaces. Real-time impedance monitoring can serve as a high-fidelity early-warning signal of electrode–tissue interface status. Because growing gas bubbles (a non-conductive phase) directly alter interface impedance, stimulation systems can detect the onset of the “electrical lensing” regime in real time. Closed-loop systems could in principle then dynamically adjust pulse parameters—to either harness bubble-assisted delivery or reduce current to avoid failure. As neural interfaces shrink to single-neuron scale, integrating such electrochemical feedback is essential to ensure chronic, high-resolution neuromodulation remains both effective and biophysically safe (*46*, *47*).

## MATERIALS AND METHODS

### Animals

Adult male C57BL/6J mice aged 14-16 weeks, weighing 25 g, were obtained from GemPharmatech and kept for 12 h in the light/12 h in the dark with sufficient food and water. All procedures were approved by the Institutional Animal Care and Use Committee (IACUC) at Laboratory Animal Center Fudan University (Permit Number: [A20250052]) and were in line with national guidelines for conducting animal experiments.

### Flexible microelectrode arrays

Flexible neural interfaces were fabricated as linear 32-channel arrays on a polyimide substrate following procedures similar to those described previously (*33*). Briefly, a sacrificial metal layer (Ni) was deposited on a glass substrate, followed by the sequential formation of a PI bottom layer, gold wires and contact pads (Pt), a PI top layer, and windows for the contact pads and channel pads using photolithography and plasma etching. Each array contained 32 platinum pads (diameter 25 µm, distance between pads 30 µm, thickness of the probe 1.3 µm) arranged in a single line (implantation length ∼1 mm). The same array was used for stimulation and recording; electrodes were connected to the printed circuit board via gold contact pads.

### 2P imaging

2P imaging was performed using a custom-built system based on a Bergamo II upright microscope (Thorlabs) equipped with a motorized XYZ stage. Excitation was provided by a femtosecond laser source operating at 1030 nm (Fibre Series, Keyun Photoelectric Technology) with a pulse duration of 180 fs and a repetition rate of 80 MHz. The laser power was controlled using a half-wave plate and a polarizing beam splitter (or simply “power control module”). Imaging was carried out using a 16× water-immersion objective (N16XLWD-PF, NA 0.80, 3.0 mm WD, Nikon). The fluorescence signal was detected via an epi-detection module (BDM1208, Thorlabs) equipped with a 525/50 nm bandpass filter and an amplified GaAsP photomultiplier tube (PMT2100R, Thorlabs). Images were acquired using galvanometer-based scanning controlled by ThorImageLS software. The field of view was 839 × 839 µm (512 × 512 pixels), resulting in a pixel size of 1.64 µm. The frame rate was set to 15.1 fps with bidirectional scanning enabled (Two-Way Alignment). Animals (n = 7) were awake and head-fixed during imaging. To visualize the vasculature, FITC-dextran (2,000 kDa) dissolved in saline (5 %, 80 µL; MaoKangBio, Shanghai, China) was injected into the retro-orbital sinus 20–30 min prior to stimulation.

### ICMS protocol and optimization

During stimulation experiments, implanted electrode in the right somatosensory cortex served as the cathode, while a stainless-steel wire in the left hemisphere functioned as the anode and ground/reference to minimize electrical noise. Prior to *in vivo* trials, stimulation parameters were optimized to maximize gas bubble formation efficiency. Based on these tests, stimulation trains were delivered at a constant frequency of 66.7 Hz with optimized burst structures (pulse train duration and number of trains) to facilitate effective electrolysis. Current amplitudes of 20, 40, 60, and 80 µA were applied while the 2P field of view was centered on arterioles and venules (diameter: 20–40 µm).

### Leakage quantification

For each stimulation event, leakage was quantified from 2P FITC–dextran images by generating a binary mask of extravascular fluorescence. Briefly, a background region of interest (ROI) was selected in the parenchyma away from vessels and the electrode track. The leakage threshold was defined as T = mean(background) + 2×SD (background). Pixels with intensity > T were classified as “leakage” and converted into a binary mask. Mask generation and area measurements were performed in Fiji using custom macros. Leakage area was computed in µm² and averaged across vessels (within animal) and then across animals. Vessels intersecting the profile were manually excluded by ROI masking prior to thresholding, so that only parenchymal regions contributed to the leakage mask.

### Electrode modification

To reduce interfacial impedance and improve stimulation stability, the working sites of the flexible electrode arrays were coated with the conductive polymer PEDOT:PSS. Electrochemical deposition was performed using a neural electrode impedance tester (NanoZ, White Matter LLC) controlled by NanoZ v1.4 software. The deposition solution consisted of 0.01 M 3,4-ethylenedioxythiophene (EDOT) and 0.1 M sodium polystyrene sulfonate (Na-PSS) (Sigma-Aldrich) dissolved in deionized water. The flexible array served as the working electrode, while a stainless-steel wire functioned as the combined reference and counter electrode. Deposition was carried out in “match impedance” mode with a target impedance of 30 kΩ. The stimulation protocol consisted of 0.05 µA current pulses with a duration of 5 s and an inter-pulse interval of 1 s. Following deposition, the arrays were rinsed and stored in a moist chamber at room temperature until implantation. Prior to use, site impedances were verified in PBS using electrochemical impedance spectroscopy (EIS, 1 Hz – 100 kHz). Only sites exhibiting an impedance of < 1 MΩ at 1 kHz were selected for experiments.

### Surgical procedure and probe implantation

Mice were anesthetized with isoflurane (2 % for induction, 1% for maintenance) in air and head-fixed in a stereotaxic apparatus (Stoelting Co., USA). Body temperature was maintained at 37 °C using a heating pad. To reduce cortical inflammation and edema, dexamethasone (1 mg/kg) was administered intraperitoneally 20 min prior to the craniotomy. A circular craniotomy (Ø 3–4 mm) was performed over the parietal bone, leaving the dura mater initially intact. Subsequently, the dura mater was carefully removed using a sharpened wire (tip diameter ∼30 µm) and fine forceps. To facilitate the insertion of the flexible electrode array, a tungsten wire shuttle (diameter: 25 µm) was used to provide stiffness. The wire was temporarily attached to the electrode shank using polyethylene glycol (PEG, MW 20,000, 5 %) as an adhesive. The electrode-shuttle assembly was mounted on the stereotaxic manipulator and inserted into the somatosensory cortex at an angle of ∼40° under visual guidance through a surgical microscope. Upon reaching the target depth (∼200-300 µm), the PEG adhesive was dissolved by applying saline solution to the insertion site. Once the shuttle detached, the tungsten wire was carefully retracted, leaving the flexible electrode array in place. The cranial window was then sealed with a circular cover glass and secured with dental cement. A custom-made metal head-plate was affixed to the skull for head fixation. Animals were allowed to recover for at least 30 days before the initiation of 2P imaging and stimulation experiments.

### Electrophysiological recording and stimulation

Extracellular neural activity and impedance were recorded using the Intan RHS Stim/Recording System (Intan Technologies, Los Angeles, CA, USA) with a sampling rate of 30 kHz (from 6 mice). The flexible electrode array was connected to an RHS headstage via a standard connector. To isolate spiking activity, the raw signal was bandpass filtered between 300 and 6000 Hz. ICMS was delivered using the integrated current stimulator of the RHS headstage. Stimulation timing was controlled by an external pulse generator (PulsePal v2, Open Ephys) via TTL triggers. The stimulation protocol consisted of 256 pulse trains delivered over a total period of 25 s. Each train had a duration of 98 ms and consisted of biphasic pulses (5 ms per phase pulse width) separated by a 5 ms inter-pulse interval. A 1 ms post-stimulation delay was applied after each train. Spike detection and sorting were performed offline using MountainSort4 software. Automated sorting results were manually curated using a custom-designed graphical user interface (GUI) to ensure cluster quality. To evaluate the impact of stimulation, the mean firing rate and spike amplitude were calculated for each channel across three-time windows: baseline (pre-stimulation), acute phase (0–1 h post-stimulation), and recovery phase (24 h post-stimulation).

### Finite Element Modeling

Numerical simulations of gas bubble growth and electric field-induced vascular permeabilization were performed using COMSOL Multiphysics 6.0 (COMSOL AB, Sweden). To simulate the expansion of the gas bubble, we employed an axisymmetric 2D model representing a platinum disk electrode (radius 20 µm) immersed in a conductive hydrogel medium (sigma = 0.7 S/m, ε_r_= 80). The Electric Currents interface was used to solve for the electric potential distribution during current stimulation pulses I(t). The time-dependent evolution of the bubble radius, R(t), was computed using a Global Ordinary Differential Equation (ODE). This equation incorporated Faraday’s law of electrolysis to calculate gas generation, balanced against opposing forces including surface tension and hydrostatic pressure.

To model vessel wall permeabilization and tracer leakage, a 2D geometry was constructed featuring a vessel cross-section (diameter 20-40 µm) embedded within the tissue volume. This model coupled the Electric Currents interface with the Transport of Diluted Species interface to simulate the convection-diffusion of a fluorescent marker (analogous to FITC-dextran). The vessel wall was defined as a boundary with voltage-dependent permeability. The permeability was governed by the local transmembrane potential ΔV_m_: it remained at baseline levels when ΔV_m_ was below a critical permeabilization threshold V_th_ and increased sharply according to a smooth step function when ΔV_m_ > V_th_. Bubble effects were incorporated phenomenologically. Rather than explicitly meshing a gas phase, we introduced a bubble-dependent coupling factor α(R) that scales the effective transmembrane potential, ΔV_m_eff_= α(R) × ΔV_m_. Here R denotes the bubble radius (constrained by the in vitro bubble–current relationship), and α(R) = (1-θ)^β^ with θ = min [1, (R/a)^2^] is the fractional coverage of the circular electrode by the bubble (i.e., the projected bubble area πR^2^ normalized by the electrode area πa^2^, capped at 1), a is the electrode radius (20 µm in our device), and β is a dimensionless exponent controlling how sharply field coupling decays with bubble coverage. Model parameters were calibrated such that the simulated dependence of leakage area on current amplitude qualitatively matched the experimental data, specifically reproducing the nonlinear transition observed between 40 µA and 60 µA. As a negative control, we evaluated a field-only variant with bubble screening disabled (α=1, R=R_min_), in which permeability depends solely on ΔV_m_. This variant produced a smooth monotonic leakage–current relationship and did not reproduce the sharp transition observed experimentally.

### Behavioral Videography and DeepLabCut Tracking

To assess motor responses in head-fixed mice (n = 7), video recordings were acquired using an infrared camera operating at 29.7 frames per second (fps). The camera was positioned frontally to capture the snout, chest, and forepaws (**Fig. 5B**). Video acquisition was synchronized with the electrical stimulation protocol. Motion analysis was performed using DeepLabCut (Python). A neural network was trained on labeled frames to track the trajectories of key anatomical landmarks: the nose, chest, and wrists of the forepaws. Movement velocity was calculated as the Euclidean distance of coordinate displacement between consecutive frames (pixels/frame). For quantitative analysis, the maximum velocity was extracted for each stimulation trial within three defined time windows: “Before” (baseline, –50 to 0 s), “Stim” (during stimulation, 0 to 25 s), and “After” (recovery, 25 to 75 s).

### Immunohistochemistry and Histological Quantification

At either 5 days or 6 months after the completion of the stimulation protocols, animals were deeply anesthetized and transcardially perfused with phosphate-buffered saline (PBS) followed by 4 % paraformaldehyde (PFA). Brains were harvested and post-fixed in PFA at 4 °C overnight. Coronal and horizontal sections (5 µm thick) were obtained using a microtome at 30 µm intervals. Sections underwent immunofluorescent staining using primary antibodies against IBA1 (microglia), GFAP (astrocytes), and cleaved caspase-3 (apoptosis marker). Visualization was performed using species-appropriate fluorophore-conjugated secondary antibodies, and nuclei were counterstained with DAPI. For quantitative analysis, two sections containing the electrode track were selected per animal (*n* = 4 mice). Using ImageJ software, the electrode track was manually segmented, and a region of interest (ROI) was defined as a concentric annulus extending 0–300 µm from the track border. The proportional area occupied by GFAP, IBA1 and cleaved Caspase-3 signals was calculated following thresholding and normalized to a contralateral ROI of equivalent geometry.

### Sample Size Variations

A total of 7 adult mice were initially enrolled in the study and successfully underwent implantation surgery, establishing the baseline cohort (**Fig. 1C**). During the initial phase at D30, all n = 7 mice successfully completed the simultaneous intravital two-photon imaging and infrared posture tracking. Following this imaging phase, one animal was excluded due to a headstage connector failure, leaving n = 6 mice to enter and complete the multi-day electrophysiological recording and sequential stimulation phase (D45–D49). Following repeated high-current stimulation, acute hardware instability and insulation failure occurred in two additional subjects. Consequently, only the final cohort of n = 4 mice successfully completed the subsequent high-current longitudinal monitoring (D65) and was processed for chronic endpoint histology later.

### Statistics

Statistical analyses were performed using OriginPro2024 and Python 3. Data are presented as mean ± SEM, unless otherwise stated. Sample size (n) refers to the number of animals unless explicitly noted otherwise (e.g., recording channels or vessels). For linear relationships (e.g., OCP vs pH, bubble area vs stimulation current), least-squares regression was applied, and the coefficient of determination (R^2^) was reported. For electrode performance and unit stability (**Fig. 4**), changes in firing rate, amplitude, and impedance were evaluated using one-sample two-tailed t-tests against a theoretical mean of zero to determine the presence of a stimulation effect. Comparisons between different stimulation intensity groups (20, 40, 60, and 80 µA) were performed using one-way ANOVA followed by Tukey’s post-hoc test for multiple comparisons. For histological quantitative analysis (**Fig. 5H**), area fractions of GFAP, Iba-1, Caspase-3 and DAPI were compared between the control (contralateral), 5-day, and 6-month groups using one-way ANOVA. Normality was assessed using the Shapiro-Wilk test; where normality could not be assumed, the Kruskal-Wallis test with Dunn’s post-hoc correction was employed. Statistical significance was defined as: *p < 0.05, **p < 0.01, and ***p < 0.001.

## Acknowledgments

The authors thank Dr. Victor Naumenko for valuable comments and critical reading of the manuscript. Large language models (including ChatGPT and Gemini) were used during the preparation of this manuscript for English language copyediting and proofreading to improve readability. Additionally, AI tools were utilized to assist in writing and optimizing the custom Python scripts used for the electrophysiological and imaging data analysis. All final text, code logic, and data interpretations were thoroughly reviewed, verified, and approved by the authors, who take full responsibility for the integrity of the content.

## Author contributions

Conceptualization: A.I., F.H.

Methodology: A.I., H.M., M.X., Ch.Y., F.H.

Investigation: A.I., H.M., F.H.

Visualization: A.I., F.H.

Supervision: F.H.

Writing—original draft: A.I.

Writing—review & editing: H.M., F.L., Zh.Ch., M.X., Ch.Y., R.L, J.W., F.H.

## Data, code, and materials availability

All data needed to evaluate and reproduce the findings presented in this paper are present in the paper and/or the Supplementary Materials. Additional data related to this paper may be requested from the authors.

