## Supplementary Materials for "Electrolytic-microbubble dynamics delineate safety thresholds during intracortical microstimulation with flexible neural interfaces"

Artem Iliasov *et al.*

**This PDF file includes:**

Supplementary Text
Figs. S1 to S5

**Other Supplementary Materials for this manuscript include the following:**
Movies S1 to S8

### Supplementary Text

Electrochemical Degradation of PEDOT: PSS Coatings Under Low-Amplitude Stimulation

Representative bright-field micrographs of a PEDOT:PSS-coated gold stimulation pad demonstrate its structural evolution before stimulation, after a single 10 μA stimulation bout, and after repeated 10 μA stimulation (Fig. S1A–C). Even at this low current amplitude, the PEDOT:PSS layer exhibits visible delamination from the underlying gold substrate, followed by lateral displacement and partial loss of the conductive coating upon repeated stimulation. These structural changes are accompanied by a progressive loss of pad integrity and precede complete electrical failure.

In contrast, platinum contacts subjected to the same stimulation protocol remain structurally intact up to substantially higher current amplitudes (see **Fig. 2E** in the main text). Real-time tracking of this degradation process is documented in **Movie S2**, which captures the delamination of the PEDOT:PSS layer during a single 10 μA stimulation. Cumulative displacement and material loss during repeated 10 μA stimulation cycles are further illustrated in **Movie S3**.

Spatiotemporal Dynamics of Microbubble Inflation and Vascular Leakage

Sequential imaging frames acquired before stimulation, during bubble growth, and following bubble deflation delineate the tight coupling between bubble dynamics and blood-brain barrier disruption (Fig. S2). While the electrochemically generated gas bubble remains attached to the vessel wall during active stimulation, extravascular FITC fluorescence remains minimal, indicating that the physical presence of the bubble temporarily restricts tracer clearance.

A pronounced leakage plume emerges immediately after bubble deflation and becomes clearly visible in the minutes following complete bubble disappearance (25 min). This extravascular FITC signal gradually dissipates and becomes indistinguishable from the background by ~32 min. The temporal profile of the fluorescence decay is highly consistent with passive diffusion and physiological clearance mechanisms rather than active, ongoing vascular leakage.

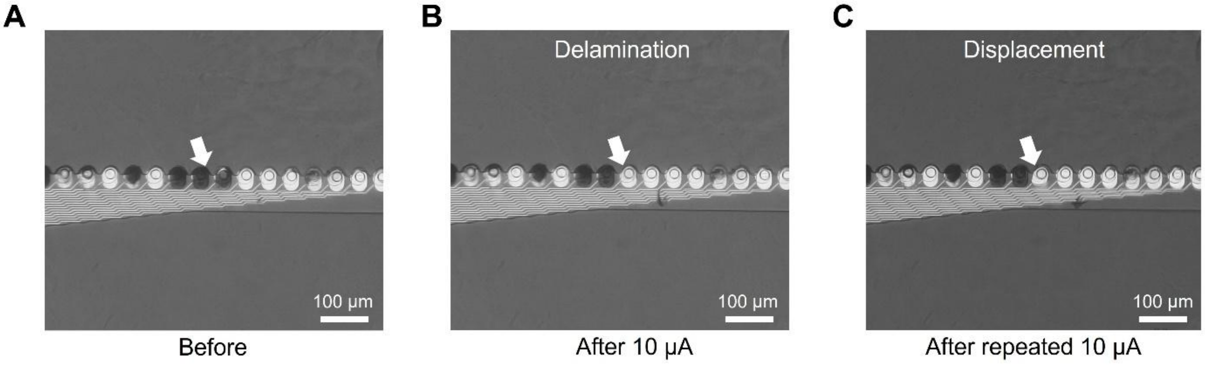


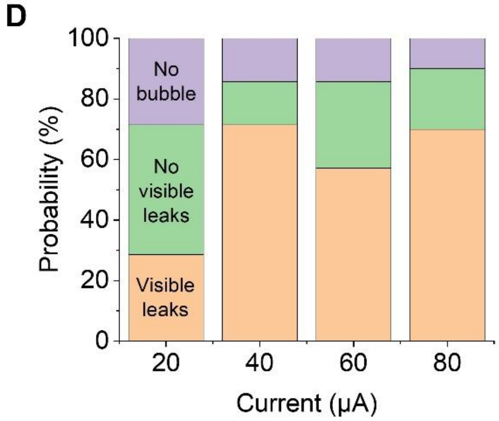


E

| Amplitude | Visible leaks | No visible leaks | No bubble | N (total) |
|---|---|---|---|---|
| 20 μA | 2 | 3 | 2 | 7 |
| 40 μA | 5 | 1 | 1 | 7 |
| 60 μA | 4 | 2 | 1 | 7 |
| 80 μA | 4 | 2 | 1 | 7 |

**Figure S1. Electrochemical degradation and bubble formation thresholds**. (**A–C**) Rapid electrochemical degradation of gold-based stimulation pads under low-amplitude stimulation. (**D–E**) Probabilistic nature of bubble formation and vascular leakage across stimulation amplitudes: (D) Stacked bar plot showing the fraction of imaging sessions resulting in three mutually exclusive outcomes for each stimulation amplitude ($N$ indicates the total number of imaging sessions at each current). (E) Tabulated counts of the same stimulation outcomes.

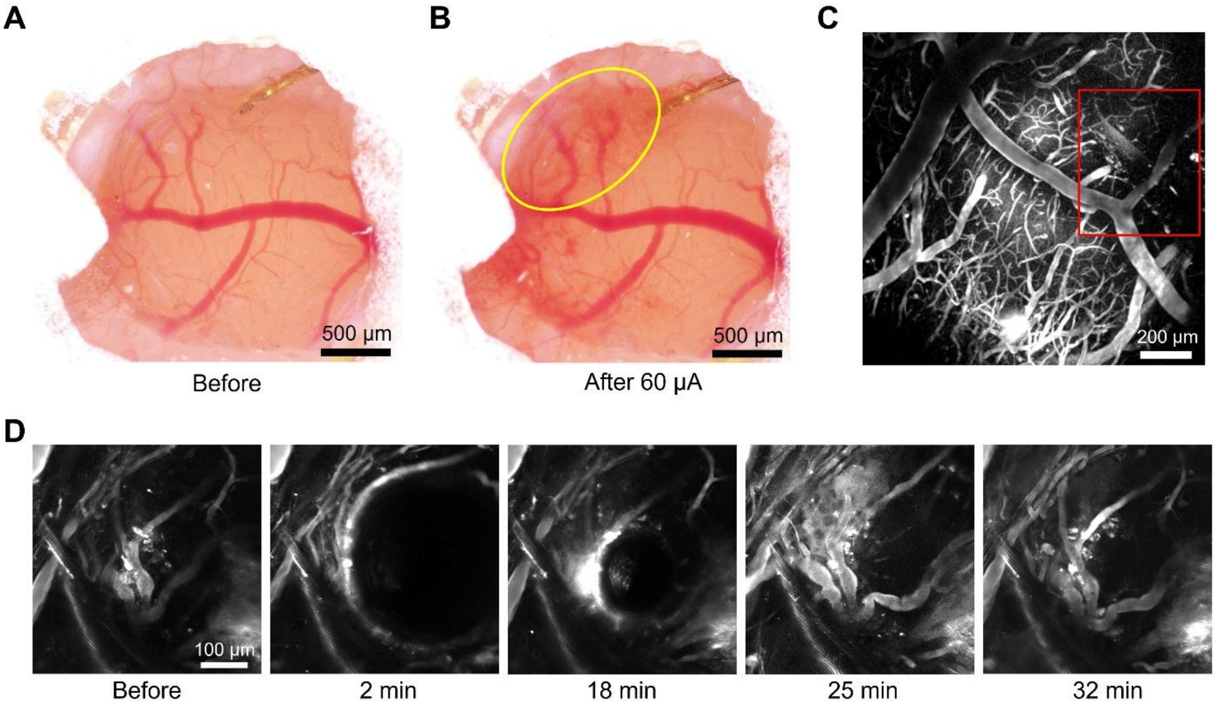


**Figure S2. Dynamics of microbubble-induced vascular leakage.** (**A, B**) Widefield visualization of large-scale vascular leakage beyond the two-photon field of view. **(C)** Overview image of the vascular network, with red boxes indicating regions shown at higher magnification below. **(D)** Sequential frames acquired before stimulation, during bubble growth, and following bubble deflation (2–32 min).

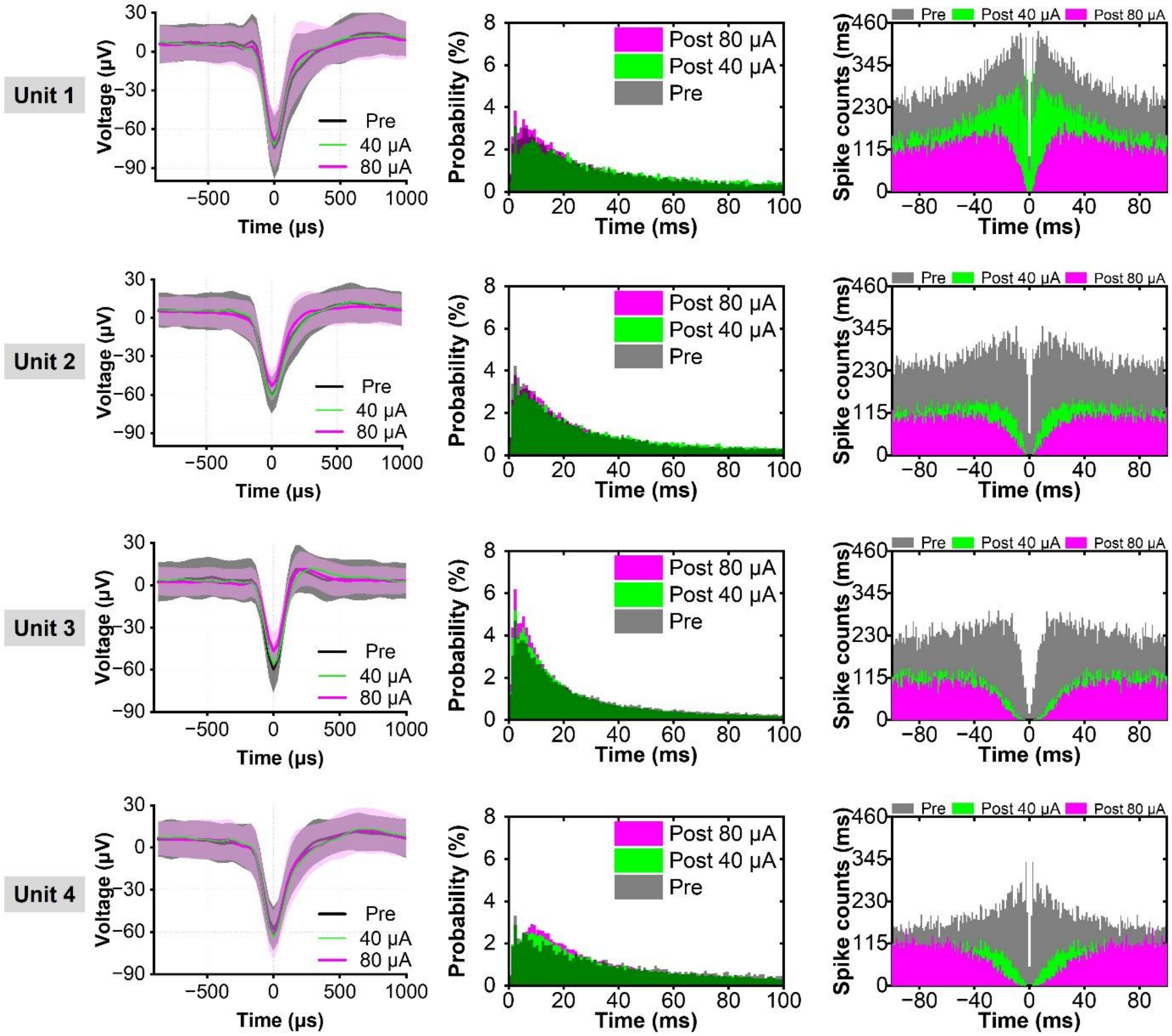


**Figure S3**. **Spike waveform stability, inter-spike interval (ISI) distributions, and autocorrelograms for additional isolated single units.**

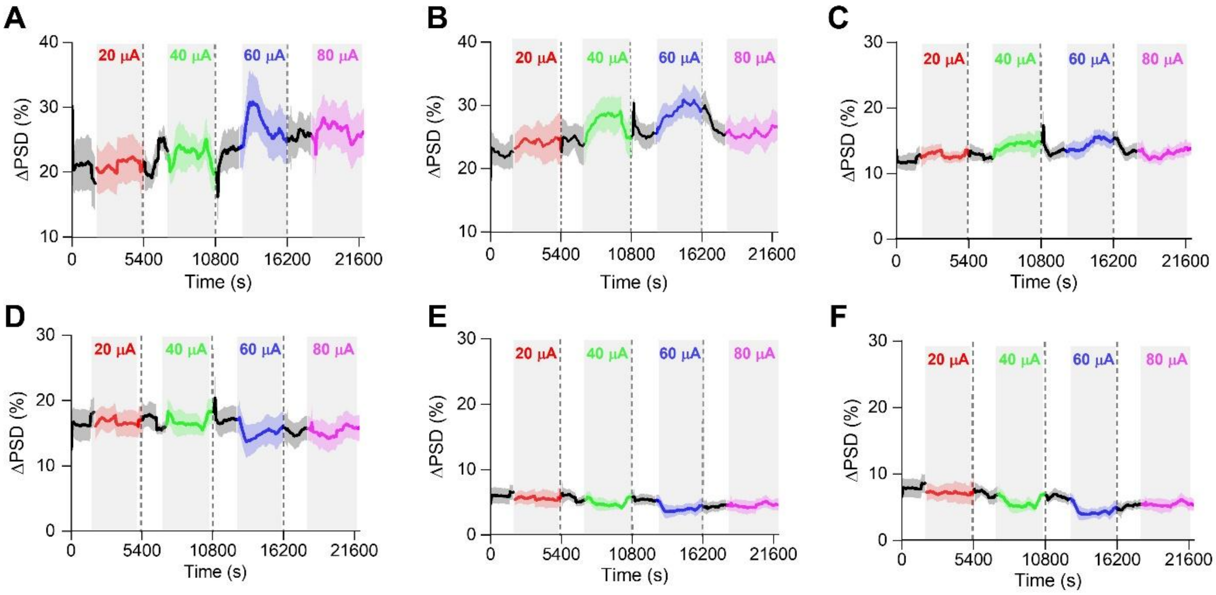


**Figure S4. Temporal dynamics of individual LFP frequency bands across different stimulation amplitudes. (A–F)** Relative changes in power spectral density (ΔPSD, %) over time (s) across consecutive stimulation epochs with increasing current amplitudes, indicated by shaded areas. **(A)** Delta, **(B)** Theta, and **(C)** Alpha frequency bands remain relatively stable and do not show consistent or significant deviations from the baseline. **(D)** Beta, **(E)** Low gamma, and **(F)** High gamma bands display amplitude-dependent post-stimulation reductions in power, most evident at intermediate and higher amplitudes (40-60 µA). Data are presented as mean (solid lines) ± SEM (shaded error bands).

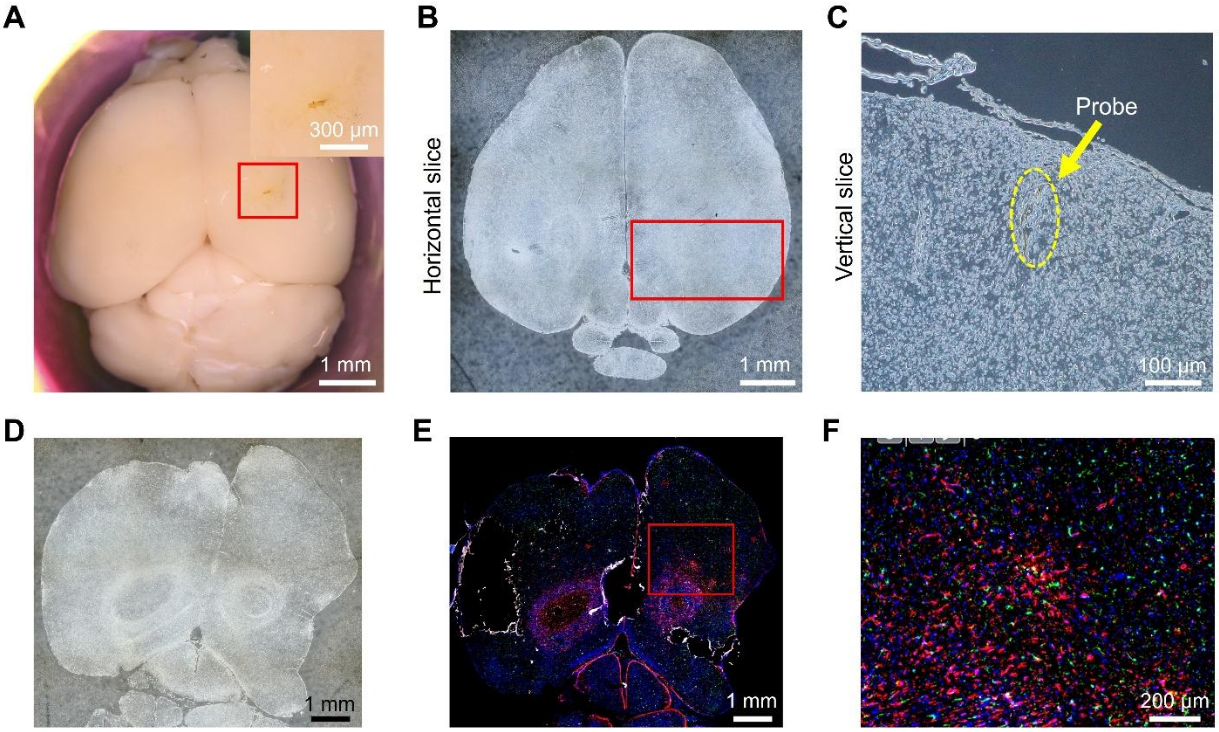


**Figure S5. Gross morphology and histological validation of implantation sites at acute and chronic time points. (A–C)** Characterization of the brain tissue corresponding to the acute immunofluorescence data presented in Fig. 5F–G. **(A)** Dorsal view of the harvested whole mouse brain, with the red boxed region indicating the electrode insertion site (inset shows a magnified view of the tissue surface. **(B)** Representative brightfield image of the horizontal brain section containing the electrode track. **(C)** High-magnification phase-contrast/brightfield micrograph showing the precise tissue track left by the microelectrode array (indicated by the yellow dashed outline). **(D–F)** Long-term biocompatibility and tissue response in a mouse brain 6 months post-stimulation (chronic time point). **(D)** Overview brightfield image of a coronal slice. **(E)** Representative low-magnification immunofluorescent coronal section showing the region of interest (red box). **(F)** High-magnification multicolor immunofluorescent image of the chronic implantation site, displaying stable tissue architecture with localized glial responses.

**Movie S1.** Gas bubble coalescence *in vitro* (1.2 % agarose gel)

A bubble nucleates at the stimulated channel and expands during the burst. Sub-visible microbubbles form along the electrode surface and rapidly coalesce into a single macroscopic bubble localized at the active site.

**Movie S2.** Real-time PEDOT:PSS delamination from a gold contact during electrical stimulation (10 µA).

**Movie S3.** Pad displacement from a gold probe during stimulation (10 µA).

**Movie S4.** Three-dimensional time-lapse reconstruction of the imaged vascular region before stimulation.

**Movie S5.** Three-dimensional time-lapse reconstruction of the same vascular region after stimulation.

**Movie S6.** Large bubble formation and erythrocyte extravasation during high-current stimulation.

**Movie S7.** Peak FITC leakage during partial bubble presence followed by gradual diffusion.

**Movie S8.** Sequential bubble formation during repeated stimulation. Vascular leakage does not occur.
